# Partial EMT and associated changes in cellular plasticity in oncovirus-positive samples

**DOI:** 10.1101/2022.08.20.504546

**Authors:** Manas Sehgal, Ritoja Ray, Joel Markus Vaz, Shrihar Kanikar, Jason A. Somarelli, Mohit Kumar Jolly

## Abstract

Oncoviruses exploit diverse host mechanisms to survive and proliferate. These adaptive strategies overlap with mechanisms employed by malignant cells during their adaptation to dynamic micro-environments and for evasion of immune attack. While the role of individual oncoviruses in mediating cancer progression has been extensively characterized, little is known about the common gene regulatory features of oncovirus-induced cancers. Here, we focus on defining the interplay between several cancer hallmarks, including Epithelial-Mesenchymal Transition (EMT), metabolic alterations, and immune evasion across major oncoviruses by examining publicly available transcriptomics data sets containing both oncovirus-positive and oncovirus-negative samples. We observe that oncovirus-positive samples display varying degrees of EMT and metabolic reprogramming. While the progression of EMT generally associated with an enriched glycolytic metabolic program and suppressed fatty acid oxidation (FAO) and oxidative phosphorylation (OXPHOS), partial EMT correlated well with glycolysis. Furthermore, oncovirus-positive samples had higher activity and/or expression levels of immune checkpoint molecules, such as PD-L1, which was associated with a partial EMT program. These analyses thus decode common pathways in oncovirus-positive samples that may be used in pinpointing new therapeutic vulnerabilities for oncovirus-associated cancer cell plasticity.

## Introduction

Oncoviruses are a class of viruses that can induce cancer in the host organism. A few examples of oncoviruses include Epstein-Barr virus (EBV), human papillomavirus (HPV), hepatitis B virus (HBV), and hepatitis C virus (HCV). These can be primarily classified as DNA tumor viruses (HPV, HBV, and EBV) and RNA tumor viruses (HCV) (McLaughlin-Drubin and Munger 2008). Such viral infections can contribute to as high as 15% of human cancers worldwide (Cao and Li 2018). Different oncoviruses are associated with specific cancer types; for example, HPV has been implicated in breast, skin, lung, cervical, and prostate cancer (McLaughlin-Drubin and Munger 2008; Xiong et al. 2017; Yin et al. 2017; Khodabandehlou et al. 2019), while HBV and HCV are strong risk factors for hepatocellular carcinoma (El-Serag 2012). While oncovirus infection does not always lead to cancer, chronic viral infections, paired with additional host factors, such as genomic instability, cell proliferation and a milieu of genetic and epigenetic modifications can ultimately initiate tumorigenesis (Tornesello et al. 2018).

A commonly observed phenomenon in cancer progression, which often associates with remodeling of the tumor microenvironment, is epithelial-mesenchymal transition (EMT). EMT is a process that involves epithelial cells losing their epithelial traits, such as cell-cell adhesion and apico-basal polarity and gaining migratory and invasive features often observed in a mesenchymal phenotype (Nieto et al. 2016). Partial or full EMT can be considered as a fulcrum of cancer cell plasticity; it is associated with multiple changes to cancer cell behavior, including migration and invasion, metabolic reprogramming, and immune evasion (Hajra et al. 2002; Yang et al. 2004; Wels et al. 2011; Dongre et al. 2017; Sciacovelli and Frezza 2017; Sahoo et al. 2021; Muralidharan et al. 2022). EMT in cancer is mediated by a host of tumor microenvironment factors, including hypoxia, matrix stiffness and crosstalk with other stromal cells (Kumar *et al*., 2014; Li *et al*., 2019; Saxena and Jolly, 2019). In addition to these factors, oncoviruses are also capable of inducing EMT in cancer cells. In 1994, Gilles and colleagues confirmed EMT and increased invasiveness in HPV33-infected cervical keratinocytes (Gilles et al. 1994). Since then, several studies have demonstrated induction of EMT by major oncoviruses (Chen et al. 2016). However, recent analyses have emphasized the spectrum of phenotypes observed across the epithelial/mesenchymal axis. Rather than a binary and complete switch to the mesenchymal state, cancer cells often display a mix of epithelial and mesenchymal traits, attaining one or more hybrid epithelial/mesenchymal (E/M) phenotypes (Tripathi et al. 2020). Thus, this more complex and nuanced understanding of epithelial-mesenchymal plasticity prompts a renewed analysis of the association between oncoviruses and EMT-like phenotypes.

Another consequence of oncovirus infection is reprogramming of host cell metabolism. Oncoviruses can rewire host cell metabolism to synthesize macromolecules important for both viral replication and tumor growth (Purdy and Luftig 2019; Mullen and Christofk 2022). Such reprogramming is observed across many oncoviruses, although the degree to which this phenomenon occurs varies based on host factors and viral requirements. This reprogramming may also contribute to promoting cancer progression (Piccaluga et al. 2018). Hence, therapeutic solutions targeting specific metabolic pathways are likely to prevent viral infections and potential oncogenesis.

Besides the association between EMT and metabolic reprogramming, oncoviruses are also observed to interact with the host immune system. Many oncoviruses, such as EBV and HPV, are associated with increased levels of PD-L1, a co-inhibitor of cytotoxic T-lymphocyte attack (Farrukh et al. 2021). Tumor cells expressing high PD-L1 are capable of evading T-cell-mediated anti-tumor responses (Han et al. 2020). Cells in partial or full EMT states often have higher levels of PD-L1 than epithelial ones (Dongre et al. 2017; Sahoo et al. 2021), suggesting that EMT can contribute to immune evasion by upregulating immune checkpoints. However, it remains to be investigated whether oncoviruses induce an increase in PD-L1 and other checkpoints, and whether this alteration in immune checkpoints is associated with a partial or full EMT.

Given the key connections between oncoviruses, EMT, metabolic plasticity, and immune evasion we sought to better understand the coupling among these axes of plasticity (EMT, metabolism, and immune evasion) in oncovirus-positive samples. To do this, we performed a comprehensive meta-analysis involving multiple human oncoviruses to analyze their associations with EMT, metabolic plasticity, and PD-L1 levels and activity – using 80 transcriptomic datasets, containing both oncovirus-positive and oncovirus-negative samples. Our meta-analysis shows that oncovirus-positive samples often associate with a partial EMT, downregulation of oxidative phosphorylation and fatty acid oxidation metabolic pathways, and an upregulation in glycolysis. Moreover, expression levels of *CD274* (gene encoding PD-L1) and activity for PD-L1 pathway, together with expression levels of immune checkpoint molecules, such as CD47, HAVCR2 and CD276, are enriched in oncovirus-positive samples. Thus, our results suggest a partial EMT phenotype and associated changes in metabolic reprogramming and T-cell mediated immune checkpoints in the oncovirus-positive samples.

## Materials and Methods

### Software and datasets

For computational and statistical analyses, Python (version 3.10) and R (version 4.1.2) were used. Microarray/RNA-sequencing datasets were downloaded from the National Center for Biotechnology Information Gene Expression Omnibus (NCBI GEO). For microarray datasets, probe-wise expression matrices were downloaded using GEOquery R Bioconductor package, and their corresponding annotation files were used to map probes to obtain the gene-wise expression. If more than one probe was mapped to the same gene, we used the average expression value of all probes mapped to the. The gene expression data was then log2-transformed (normalized). For RNA-seq datasets, sample-wise raw counts for each dataset (denoted by GSE ID) were downloaded from NCBI GEO database, normalized for gene length, and transformed to TPM (transcripts-per-million) values which were then log2-transformed to obtain the final expression values for each gene per sample.

### EMT, PD-L1, and Metabolic scoring metrics

Epithelial-mesenchymal transition (EMT) scores were generated using KS and 76GS scores (Chakraborty et al. 2020; Mandal et al. 2022), and single-sample Gene Set Enrichment Analysis (ssGSEA). PD-L1 activity scores were also obtained using ssGSEA. The KS score uses a two-sample Kolmogorov–Smirnov (KS) test to quantify the E/M (Epithelial/ Mesenchymal) status of a given sample. This scoring metric employs 218 gene and 315 gene signatures for tumor samples and cell line samples, respectively. To acquire the EMT score, we obtain cumulative distribution functions (CDFs) for the two signatures (Epithelial and Mesenchymal). The maximum distance between these CDFs is utilized as the test statistic for a two-sample KS test giving a final E/M score in the range of [-1, 1]. Positive and negative scores denote mesenchymal and epithelial phenotypes, respectively. The 76GS method utilizes 76 gene signatures to calculate E/M scores for each sample. The score of each sample is subtracted from the mean of all samples such that the resulting mean score is zero. The negative scores indicate a mesenchymal phenotype, whereas positive scores represent an epithelial phenotype. The 76GS score does not have a defined range.

### Single-sample Gene Set Enrichment Analysis

Single-sample Gene Set Enrichment Analysis (ssGSEA) assigns an enrichment score for each sample pairing with a given gene set which signifies the degree of enrichment of that set of genes from a given pathway toward the top or bottom of an input list. It utilizes a preprocessed sample-wise gene expression matrix as input and generates scores using a distinct algorithm (Aravind et al. 2005). To obtain ssGSEA scores for different pathways, we obtained hallmark genesets (as described below) for different pathways from the MSigDB repository and calculated the scores for each sample using the GSEAPY python library.

### Genesets used for analysis

Genesets for hallmark EMT, FAO, OXPHOS, glycolysis, HIF-1α and PD-L1 were obtained from MSigDB. Genes used to obtain KS scores previously were used as genesets for calculating the KS epithelial and mesenchymal scores separately (referred as ‘Epi’ and ‘Mes’ scores) **(Table S7**). ssGSEA scores were calculated in the GSEAPY python library for all datasets using these hallmark genesets to obtain normalized enrichment scores (NES) for each pathway across all samples.

### Correlation analysis

Correlation between scores obtained from different EMT scoring metrics and between EMT scores and ssGSEA scores for metabolism and PD-L1 were calculated using Pearson’s correlation. A two-tailed Student’s t-test with unequal variance was performed to determine the statistical reliability of the observations *(p<0*.*05)*. Only those datasets were considered whose correlation coefficient (R) was less than −0.3 (significant negative correlation) or greater than 0.3 (significant positive correlation). Similar parameters were set as cut-off for the 2D correlation plots. Comparison between metrics and generation of plots and figures was carried out using R (version 4.1.2).

### Comparison between oncovirus positive and negative samples

To check if oncovirus-positive samples were enriched in a pathway, the mean score of enrichment of each pathway in oncovirus-positive samples was compared with that of oncovirus-negative samples for each dataset. Datasets showing a significant difference between the means are analyzed *(p<0*.*05)*.

### Probability plots

For quantifying the probability of a dataset correlating significantly (p<0.05), either positively (r>0.3) or negatively (r<-0.3) for a given pair of metrics in the expected direction of association, the probability was calculated as the ratio of the number of datasets showing significant correlation in the expected direction (positive correlation for Mes vs. KS and Epi vs. 76GS, negative correlation for Mes vs. 76GS and Epi vs. KS) to the total number of datasets showing significant correlation between those two metrics in either direction. The higher this probability value, the better the compliance between the two corresponding EMT scoring metrics.

### Venn Diagrams

To assess the degree of overlaps/consistency between each pairwise comparison of EMT scoring metrics as well as between pairs of EMT and metabolic scores applied to our set of 146 datasets, four-way Venn diagrams were plotted. Each of the four sets in the Venn diagrams have a total number equal to the datasets showing a correlation between any two concerned comparisons in the expected direction.

## Results

### Oncovirus-positive samples associate with a partial EMT phenotype

To quantify the extent of EMT, we used four distinct transcriptomics-based EMT scoring metrics. Two of those metrics are KS (Tan et al. 2014) and 76GS (Byers et al. 2013) scores. Both these metrics use distinct sets of gene lists corresponding to epithelial and mesenchymal phenotypes and generate a score to identify the position of samples along the epithelial-hybrid-mesenchymal spectrum (Chakraborty et al. 2020; Mandal et al. 2022). EMT is a non-linear process where cells can take multiple trajectories in the high-dimensional gene expression space, and the loss of epithelial and gain of mesenchymal traits need not always be completed coupled, as is often tacitly assumed (Watanabe et al. 2019). Given these features of EMT, we also quantified the enrichment of epithelial (Epi) and mesenchymal (Mes) signatures separately using single-sample Gene Set Enrichment Analysis (ssGSEA) scores for KS epithelial and KS mesenchymal gene lists, respectively.

A higher KS or Mes score signifies a more mesenchymal phenotype, whereas a higher 76GS or Epi score signifies a more epithelial state. Thus, as expected, out of the 80 datasets used in this meta-analysis, 53 (66.25%) datasets showed a strong positive correlation between Epi and 76GS scores (**Fig 1A**), and 57 (71.25%) datasets displayed a positive association between Mes and KS scores (**Fig 1B**). Consistent with these results, KS and Epi scores were significantly negatively correlated with one another in 43 (53.7%) datasets, and positively correlated in only 4 (5%) datasets (**Fig 1C**). Similarly, KS and 76GS scores were negatively correlated with one another in 41 (51.2%) datasets, and positively correlated in just 6 (7.5%) datasets (**Fig 1D**).

**Figure 1.**
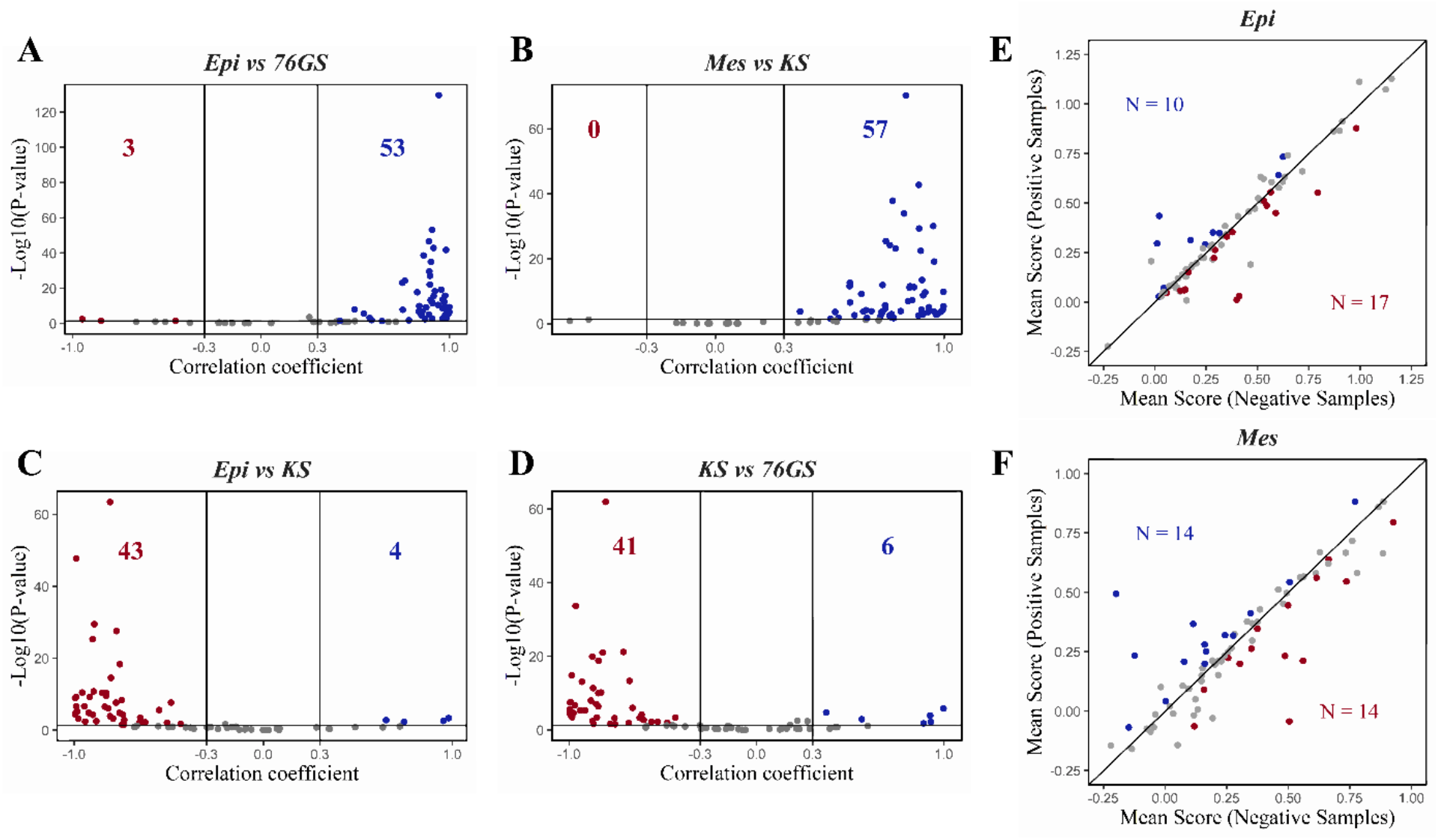
Different scoring metrics quantify E/M phenotypes in datasets containing oncovirus-infected samples. **A)** Volcano plot illustrating the Pearson correlation coefficient (x-axis) and the −log10(p-value) (y-axis) for Epi vs. 76GS scores. Vertical boundaries are set at correlation coefficients corresponding to 0.3 and -0.3 and the cut-off for significant correlation is set at p<0.05. Same as A) but for **B)** Mes vs. KS scores, **C)** Epi vs. KS and **D)** KS vs. 76GS scores. **E)** Scatter plot depicting mean Epi scores for oncovirus-positive (y-axis) and oncovirus-negative samples (x-axis) across datasets. ‘N’ indicates the number of datasets with a significant difference between the two mean scores (p<0.05). The datasets with higher mean scores for positive samples are displayed as blue datapoints, while the datasets with higher mean scores for negative samples are shown in red. Datasets with no significant difference in scores for positive vs. negative samples are shown in gray. **F)** Same as E) but for Mes scores.

Further, the Mes scores predominantly correlated negatively with both the Epi and 76GS scores (**Fig S1A-B**). Among all the above-mentioned six pairwise comparisons, the (Mes, KS) pairwise correlation had the maximum probability of being associated in the expected direction of association, followed by the (Epi, 76GS) pairwise association (**Fig S1C**). Moreover, the four-way Venn diagram revealed that 31 datasets showed complete consistency among all four pairwise comparisons, i.e., KS scores are negatively correlated with Epi scores, but positively with Mes scores, and 76GS scores are positively correlated with Epi scores, but negatively with Mes scores **(Fig S1D)**. Collectively, these observations highlight the consistency between these scoring metrics in terms of analyzing EMT-related changes in these datasets containing oncovirus-positive and negative samples.

Next, using these metrics, we investigated whether there was a difference in the epithelial and/or mesenchymal status of the oncovirus-positive vs. that of oncovirus-negative samples. For each dataset, we calculated the mean Epi score and the mean Mes score for all oncovirus-positive and oncovirus-negative samples. In 27 out of 80 datasets, we noticed a significant (p < 0.05) difference in the mean Epi scores between oncovirus-positive and –negative samples. In 17 out of those 27 datasets (63%), oncovirus-negative samples had a higher Epi score than that of oncovirus-positive samples, suggesting that the epithelial gene signature can be downregulated in the presence of an oncovirus infection. For the Mes geneset, 28 datasets showed a significant difference *(p<0*.*05)* in the mean scores, and in 14 of those datasets (50%), oncovirus-positive samples showed a higher Mes score than the oncovirus-negative ones, while in the remaining 14 (50%) datasets, an opposite trend was observed (**Fig 1F**). Thus, this analysis suggests that the oncovirus-positive samples do not have an enriched mesenchymal status as compared to oncovirus-negative samples. The features of a downregulated epithelial signature with an unaltered mesenchymal signature suggests a partial EMT or hybrid E/M phenotype of oncovirus-positive samples.

### Oncovirus-positive samples have reduced OXPHOS and FAO activity levels

Our previous pan-cancer meta-analysis for 180 datasets revealed that EMT was associated with a decreased activity of oxidative phosphorylation (OXPHOS) and fatty acid oxidation (FAO) pathways (Muralidharan et al. 2022). We capitalized on this analysis to dissect differences in OXPHOS and FAO levels for oncovirus-positive vs. oncovirus-negative samples. To do this, we first probed whether the earlier observed association between EMT and metabolic reprogramming was found in oncovirus-infected samples and quantified differences in metabolic activity of the oncovirus-positive vs. oncovirus-negative samples.

First, we correlated the ssGSEA activity scores of hallmark metabolic pathways – fatty acid oxidation (FAO) and oxidative phosphorylation (OXPHOS) – with EMT scoring metrics across datasets. FAO predominantly correlated positively with an epithelial gene signature and negatively with a mesenchymal gene signature. A total of 25 out of 80 datasets had positive correlations between FAO and Epi scores, with 8 datasets showing an opposite trend. Similarly, FAO and Mes scores correlated negatively in 21 datasets and displayed the reverse trend in only 8 datasets (**Fig 2A**). Consistent results were observed when correlating FAO enrichment with the 76GS and KS scores (**Fig S2A**). Together, these results are reminiscent of our previous observations (Muralidharan et al. 2022) and consistent with reports that virus infection leads to a shift from fatty acid oxidation to fatty acid synthesis (Sumbria et al. 2021).

**Figure 2.**
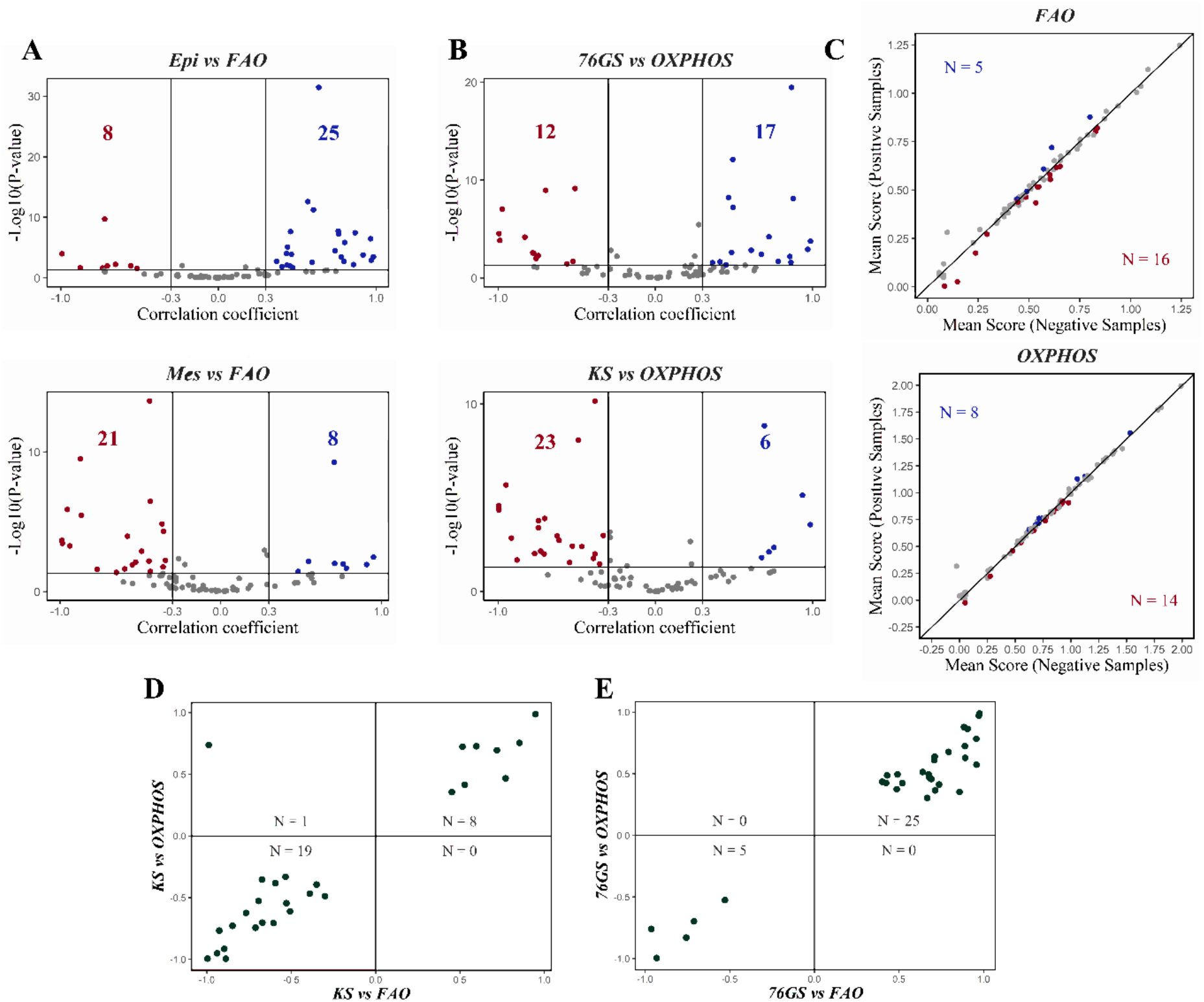
Association of EMT with OXPHOS and FAO programs in oncovirus-positive samples. **A)** Volcano plot illustrating the Pearson correlation coefficient (x-axis) and the −log10(p-value) (y-axis) for Epi vs. FAO scores (top) and Mes vs. FAO scores (bottom). Vertical boundaries are set at correlation coefficients corresponding to 0.3 and -0.3, and the cut-off for significant correlation is set at p<0.05. **B)** Same as A) but for 76GS vs. OXPHOS scores (top) and KS vs. OXPHOS scores (bottom). **C)** Scatter plots depicting mean FAO (top) and OXPHOS scores (bottom) for oncovirus-positive (y-axis) and negative samples (x-axis) across datasets. ‘N’ indicates the number of datasets with a significant difference between the two mean scores (p<0.05). The datasets with higher mean scores for positive samples are displayed as blue datapoints, while the datasets with higher mean scores for negative samples are shown in red. The datasets showing no significant difference (p>0.05) in mean scores for positive and negative samples are shown in gray. **D)** 2D scatter plot illustrating correlation coefficients of KS vs. FAO scores (x-axis) and KS vs. OXPHOS (y-axis). ‘N’ indicates the number of datasets that lie in the respective quadrant. **E)** Same as D) but for 76GS vs. FAO scores (x-axis) and 76GS vs. OXPHOS (y-axis).

Similar to the trends observed for FAO, OXPHOS also correlated positively with an epithelial state in a majority of data sets. Upon correlating OXPHOS with EMT scores, we found that OXPHOS primarily correlated negatively with a mesenchymal program and positively with an epithelial program – 23 datasets displayed a negative correlation between OXPHOS and KS scores, while only 6 datasets showed the reverse trend (**Fig 2B**, bottom). Similar behavior was evident when using Mes scores (**Fig S2B**). On the other hand, the relationship between OXPHOS and epithelial scores was more ambiguous. While 17 datasets showed a positive correlation of OXPHOS with 76GS, 12 datasets displayed a negative correlation. Similarly, in 14 datasets, OXPHOS correlated positively with an Epi score, but in 15 datasets, an opposite trend was observed (**Fig 2B**, top; **Fig S2B**). Together, these analyses suggest that OXPHOS is more likely to correlate negatively with a mesenchymal phenotype (i.e., KS and Mes scores) in the context of oncovirus-infected samples.

Next, we quantified the mean OXPHOS and FAO scores to interrogate changes in metabolic reprogramming in oncovirus-positive and -negative samples. A total of 21 out of 80 datasets showed a significant (p<0.05) difference in mean FAO scores for oncovirus-positive and oncovirus-negative samples. Only 5 out of 21 datasets (23.8%) had a higher FAO score for oncovirus-positive samples as compared to -negative ones, while in 16 datasets (76.2%), oncovirus-negative samples had a higher FAO pathway enrichment score. Similar results were found in the case of OXPHOS, wherein 14 out of 22 datasets (63.6%) had a higher OXPHOS score for oncovirus-negative samples (**Fig 2C**). These analyses suggest that both FAO and OXPHOS are often downregulated in oncovirus-positive samples.

We also examined the pairwise associations between OXPHOS and FAO to interrogate if some combinations of association were more predominant than others in the context of their correlation with EMT. For instance, in 28 datasets, KS scores were significantly correlated with both FAO and OXPHOS scores either positively or negatively. Among these 28 datasets, in 19 datasets (67.9%), the KS scores correlated negatively with both FAO and OXPHOS; in 8 datasets (29.6%), FAO and OXPHOS both correlated positively with KS scores (**Fig 2D**). Similar trends were observed when using other EMT scoring metrics – 76GS (**Fig 2E**), Epi and Mes (**Fig S2C**). FAO and OXPHOS scores correlated either positively or negatively with 76GS scores in 30 datasets. In 25 out of the 30 datasets (83.3%), 76GS scores correlated positively with both FAO and OXPHOS, while in 5 datasets (16.7%), they correlated negatively (**Fig 2E**).

Together, these results suggest that oncovirus-positive samples tend to have reduced OXPHOS and FAO activity, a pattern consistent with their partial EMT state and with our previous pan-cancer observations linking partial EMT with alterations in metabolism (Muralidharan et al. 2022). Interestingly, the association between EMT and FAO is stronger than that for EMT with OXPHOS in the context of oncovirus infection.

### Oncovirus-positive samples exhibit enrichment of glycolysis

After investigating the changes in OXPHOS and FAO levels, we tested for changes in glycolysis levels. Glycolysis is often upregulated in cancer cells (Warburg effect) to compensate for an increased ATP demand, proliferation, and survival (Shiraishi et al. 2015). To analyze the association of glycolysis with the process of EMT, we calculated ssGSEA enrichment scores for the hallmark glycolysis gene set and correlated the ssGSEA scores with different EMT scoring metrics across the 80 datasets.

We observed that glycolysis correlated positively with both epithelial and mesenchymal signatures (**Fig 3A, S3A**). This trend seems unexpected according to the canonical EMT paradigm in which epithelial and mesenchymal programs are thought to be strongly antagonistic to one another and expected to associate with diverse phenomena in opposite directions. However, EMT is a multi-dimensional process in which downregulation of an epithelial program is not necessarily strongly connected to the upregulation of a mesenchymal program and *vice versa*, thereby allowing multiple trajectories in terms of molecular and functional changes (Sahoo et al. 2022). From this perspective, it is perhaps not surprising that a positive association exists between glycolysis and both epithelial and mesenchymal programs (**Fig 3A**, top: 31 out of 36 datasets show positive correlation with epithelial; **Fig 3A**, bottom: 23 out of 29 datasets show positive correlation with mesenchymal). To further interrogate these relationships, we analyzed the relationship between epithelial and mesenchymal scores with HIF1α, a known mediator of the glycolytic pathway. Consistent with the relationships between glycolysis and EMT scores, the HIF1α pathway correlated positively with both an epithelial and a mesenchymal program. In 23 out of 29 datasets (79.3%), HIF1α correlated positively with the epithelial signature while it correlated positively with the mesenchymal signature in 28 out of 30 datasets (93.3%) (**Fig 3B**). Similar relationships existed between HIF1α and 76GS and KS scores (**Fig S3B)**. In 16 out of 27 (59.3%) datasets, glycolysis activity levels showed a higher score for oncovirus-positive samples as compared to oncovirus-negative samples. For HIF1α activity levels, 66.6% of datasets (18 out of 27) had a higher HIF1α activity score for oncovirus-positive samples (**Fig 3C)**. These trends point towards glycolysis upregulation in the presence of an oncovirus infection.

**Figure 3.**
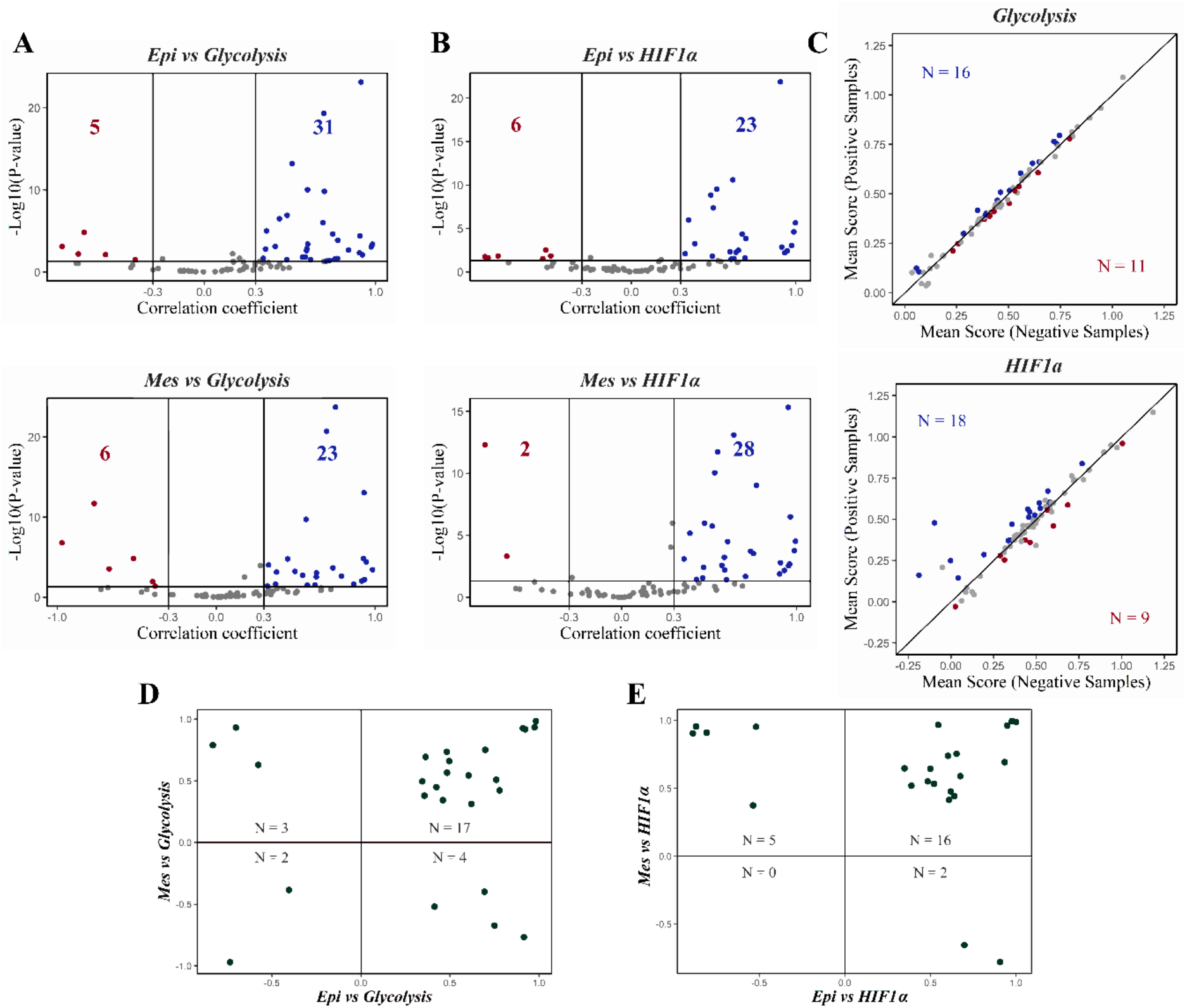
Association of glycolysis activity level (and its master regulator HIF1α) with partial EMT and oncovirus infection. **A)** Volcano plot illustrating the Pearson correlation coefficient (x-axis) and the −log10(p-value) (y-axis) for Epi vs. Glycolysis scores (top) and Mes vs. Glycolysis scores (bottom). Vertical boundaries are set at correlation coefficients corresponding to 0.3 and -0.3, and the cut-off for significant correlation is set at p<0.05. **B)** Same as A) but for Epi vs. HIF1α (top) and Mes vs. HIF1α scores (bottom). **C)** Scatter plots depicting mean FAO (top) and OXPHOS scores (bottom) for oncovirus-positive (y-axis) and negative samples (x-axis) across datasets. ‘N’ indicates the number of datasets with a significant difference between the two mean scores (p<0.05). The datasets with higher mean scores for positive samples are displayed as blue datapoints, while the datasets with higher mean scores for negative samples are shown in red. The datasets showing no significant difference (p>0.05) in mean scores for positive and negative samples are shown in gray. **D)** 2D scatter plot illustrating correlation coefficients of Epi vs. Glycolysis scores (x-axis) and Mes vs. Glycolysis (y-axis). ‘N’ indicates the number of datasets that lie in the respective quadrant. **E)** Same as D) but for Epi vs. HIF1α scores (x-axis) and Mes vs. HIF1α (y-axis).

We next asked whether the glycolysis pathway gene set correlated positively with epithelial and mesenchymal programs in the same dataset(s). Pairwise correlations between glycolysis and epithelial and mesenchymal gene signatures indicated that glycolysis correlates positively with both Epi and Mes scores (17/26 = 65.4% datasets) **(Fig 3D**). Similar association was observed in the case of HIF1α, with HIF1α correlating positively with both the epithelial and mesenchymal programs in 16 out of 23 datasets (69.6%) (**Fig 3E**). Overall, the positive association of glycolysis and its major driver, HIF1α, with both epithelial and mesenchymal gene signatures suggests that glycolysis may be correlated with a partial EMT program. This association may underlie the observations about enrichment of glycolysis in oncovirus-positive samples (**Fig 3C**), given the enrichment of partial EMT status in these samples (**Fig 1E-F**).

### Enrichment of PD-L1 signature in oncovirus-positive samples and associated changes in EMT and metabolic reprogramming

Oncoviruses such as EBV and HPV have been associated with increased PD-L1 levels (Farrukh et al. 2021). Likewise, EMT is also correlated with upregulation of immune checkpoints (Ware et al. 2020; Chakraborty et al. 2021). These relationships prompted us to investigate whether oncovirus-positive samples were enriched in PD-L1 activity and/or expression. Out of 26 datasets that showed a difference in mean PD-L1 activity levels for oncovirus-positive vs. oncovirus-negative samples, 18 (69.2%) datasets showed a higher PD-L1 activity score for oncovirus-positive samples as compared to oncovirus-negative samples (**Fig 4A**, top). Similar trends were noticed for expression levels of *CD274* (gene encoding for PD-L1) (**Fig 4A**, bottom). A total of 12/21 (57.1%) datasets displayed significant *(p<0*.*05)* upregulation of *CD274* mRNA levels for oncovirus-positive samples as compared to oncovirus-negative samples. Overall, this enrichment of the PD-L1 pathway signature in oncovirus-positive samples suggests a possible relationship between oncovirus infection and upregulation of immune checkpoints like PD-L1 and its associated gene, *CD274*.

**Figure 4.**
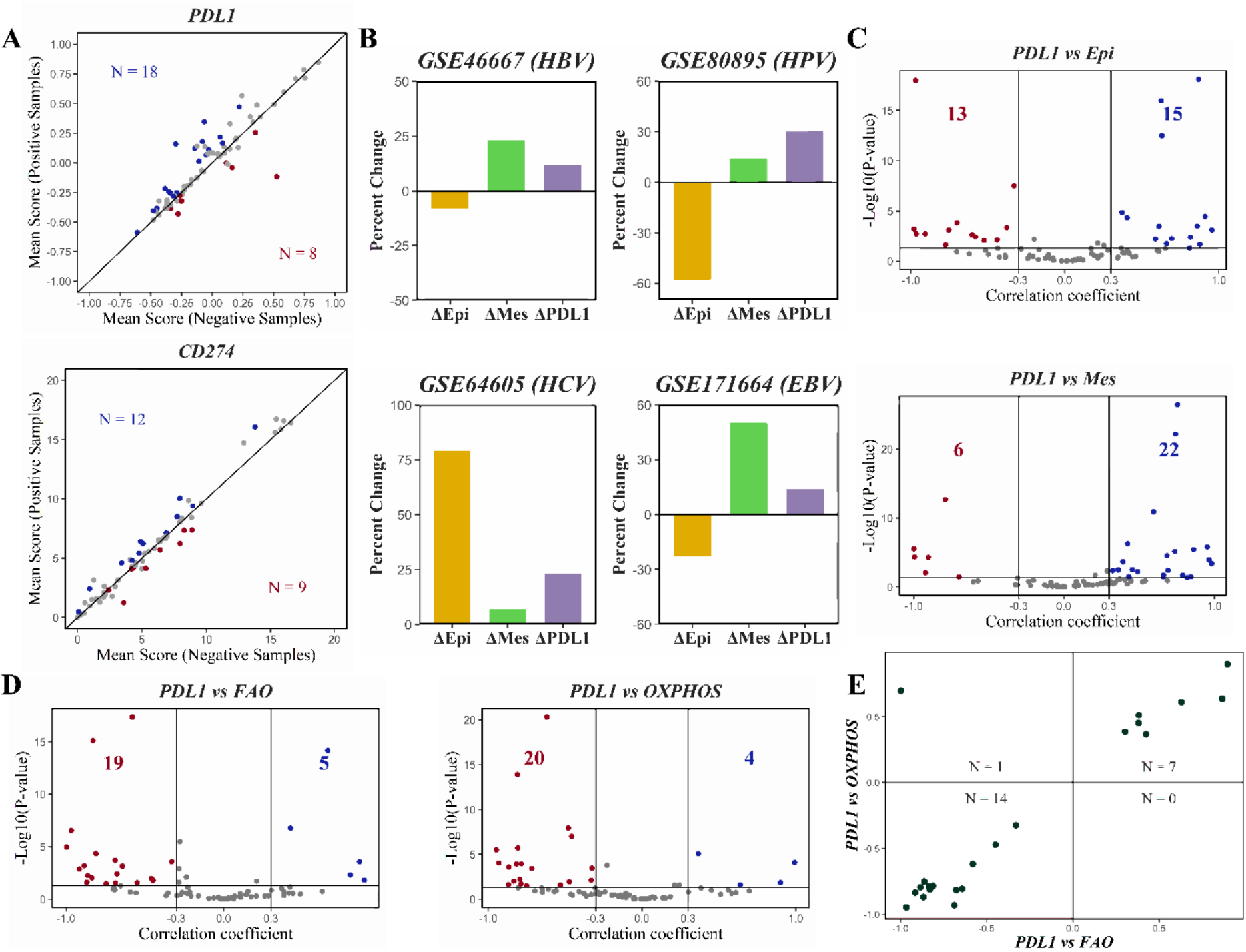
Oncovirus induced EMT-related changes in PD-L1 and different metabolic axes. **A)** Scatter plots depicting mean PD-L1 scores (top) and CD274 gene expression values (bottom) for oncovirus-positive (y-axis) and negative samples (x-axis) across datasets. ‘N’ indicates the number of datasets with a significant difference between the two mean scores (p<0.05). The datasets with higher mean scores for positive samples are displayed as blue datapoints, while the datasets with higher mean scores for negative samples are shown in red. The datasets showing no significant difference (p>0.05) in mean scores for positive and negative samples are shown in gray. **B)** Bar plots depicting relative percentage change (of positive samples with respect to negative samples) in Epi (yellow), Mes (green), and PD-L1 (purple) scores for particular datasets (denoted by GSE IDs). **C)** Volcano plots illustrating the Pearson correlation coefficient (x-axis) and the −log10(p-value) (y-axis) for PD-L1 vs. Epi score (top) and PD-L1 vs. Mes score (bottom). Vertical boundaries are set at correlation coefficients corresponding to 0.3 and -0.3, and the cut-off for significant correlation is set at p<0.05. **D)** Same as C) but for PD-L1 vs. FAO scores (left), PD-L1 vs. OXPHOS scores (right). **E)** 2D scatter plot illustrating correlation coefficients of PD-L1 vs. FAO (x-axis) and PD-L1 vs. OXPHOS scores (y-axis). ‘N’ indicates the number of datasets that lie in the respective quadrant.

After examining the association of oncovirus infection in samples with enrichment of the PD-L1 pathway gene set across datasets, we investigated the trends between changes in levels of Epi, Mes scores and activity of the PD-L1 geneset simultaneously. Here, we discuss four representative datasets, one each for a different oncovirus, and we noticed that while PD-L1 activity and Mes scores increase in all cases to varying degrees when comparing oncovirus-positive and oncovirus-negative samples. Similarly, with the exception of HCV, Epi scores are lower for oncovirus-positive samples as compared to oncovirus-negative samples (**Fig 4B**). Together, these observations suggest that the changes in Epi, Mes, and PD-L1 activity scores can be coordinated and thus observed concomitantly.

Given the relationship between PD-L1 and EMT (Ware et al. 2020; Chakraborty et al. 2021), we also analyzed the correlations between PD-L1 activity and EMT scores across our entire cohort of datasets. We observed that the PD-L1 activity score was more likely to be positively correlated with a Mes score (22 out of 28 datasets in **Fig 4C**) than with an Epi signature (15 out of 28 datasets in **Fig 4C**). Furthermore, a similar trend was observed for *CD274* mRNA levels versus Epi and Mes scores (**Fig S4D-E**). These analyses suggest that enrichment of PD-L1 scores is more aligned with the presence of a mesenchymal signature rather than the absence of an epithelial one. This trend highlights a potential association of PD-L1 with a partial EMT program as noted experimentally (Aggarwal et al. 2021; Dongre et al. 2021; Sahoo et al. 2021) in which a loss of epithelial traits and gain of mesenchymal ones can be considered as semi-independent properties (Sahoo et al. 2022). Moreover, upon plotting the pairwise association of PD-L1 with FAO and OXPHOS, the predominant association observed was that a PD-L1 signature correlated negatively with both the pathways, as observed in 63.6% of datasets (**Fig 4D-E**). Interestingly, PD-L1 activity was also correlated negatively with glycolysis (**Fig S4A**). Overall, these analyses suggest that the PD-L1 activity signature is likely to be associated negatively with metabolic reprogramming in the context of oncovirus-infected samples.

### Oncovirus infection is associated with the upregulation of immune checkpoint markers

We also determined the association between the presence of oncovirus infection and mRNA expression levels of additional immune checkpoint markers, including *CD276* (encodes B7-H3), *CD47* (encodes Cluster of Differentiation 47), *HAVCR2* (encodes TIM-3) and *LGALS9* (encodes galectin 9) (Liu et al. 2015; Holderried et al. 2019; Yang et al. 2020). Similar to trends seen for CD274, oncovirus-positive samples showed a significant upregulation in mRNA levels of *CD47* (78.2% of datasets), *HAVCR2* (64.7%) and *LGALS9* (71.4%) as compared to oncovirus-negative ones (**Fig 5A**). However, *CD276* did not show any such enrichment.

**Figure 5:**
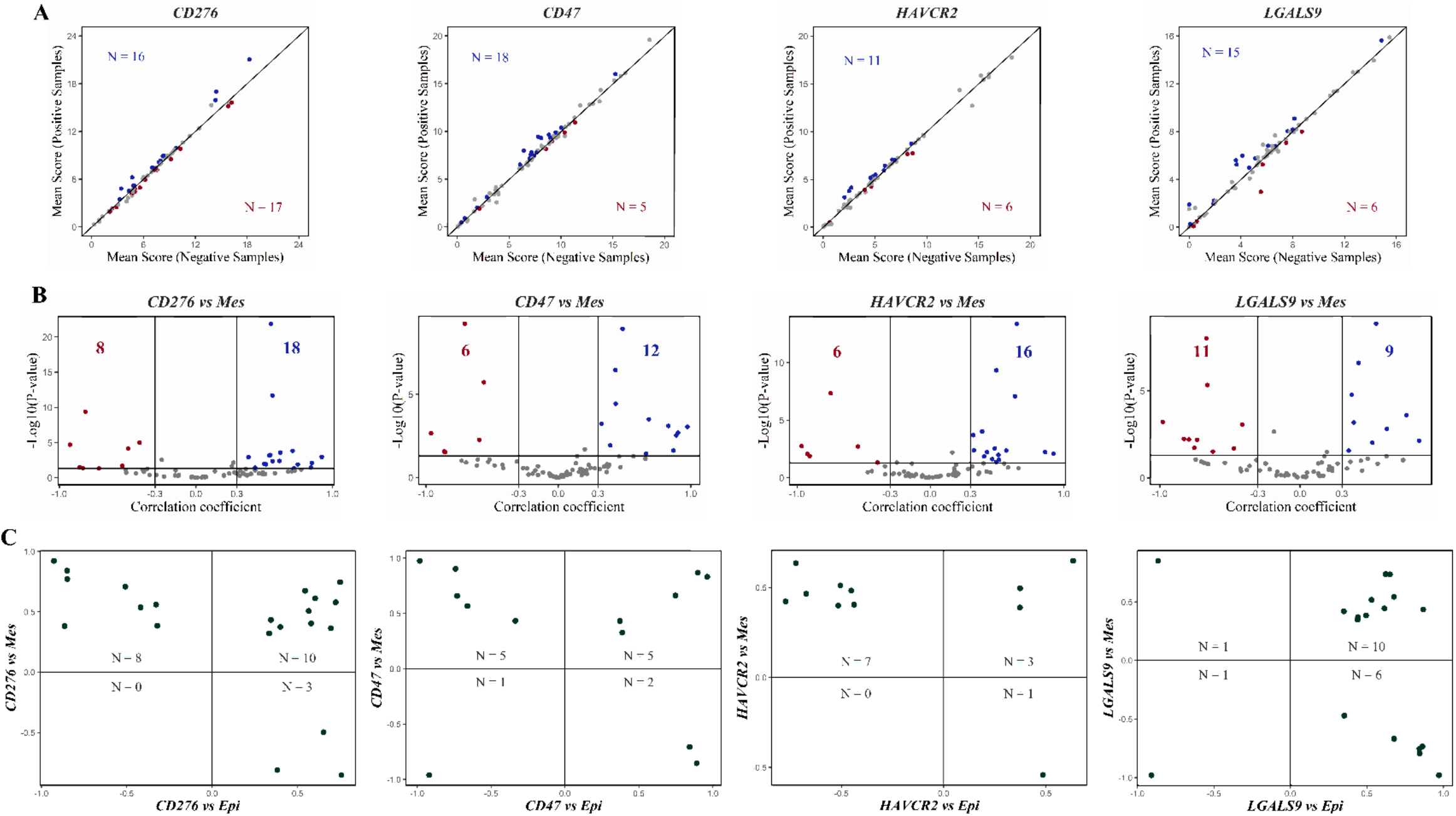
Association of gene expression of different immune checkpoints with EMT and oncovirus infection. **A)** Scatter plots depicting the mean expression values for oncovirus-positive (y-axis) and negative samples (x-axis) across datasets: (from left to right) CD276, CD47, HAVCR2 and LGALS9. ‘N’ indicates the number of datasets with a significant difference between the two mean scores (p<0.05). Datasets with higher mean scores for positive samples are displayed as blue datapoints, while those with a higher mean scores for negative samples are shown in red. Datasets showing no significant difference (p>0.05) in mean scores for positive and negative samples are shown in gray. **B)** Volcano plots illustrating the Pearson correlation coefficient (x-axis) and the −log_10_(p-value) (y-axis) for CD276 vs. Mes (left), CD47 vs. Mes (middle-left), HAVCR2 vs. Mes (middle-right) and LGALS9 vs. Mes scores (right). Vertical boundaries are set at correlation coefficients corresponding to 0.3 and -0.3, and p<0.05. **C)** 2D scatter plot illustrating correlation coefficients of CD276 vs. Epi scores (x-axis) & CD276 vs. Mes (y-axis) (left), CD47 vs. Epi scores (x-axis) & CD47 vs. Mes (y-axis) (middle-left), HAVCR2 vs. Epi scores (x-axis) & HAVCR2 vs. Mes (y-axis) (middle-right) and HAVCR2 vs. Epi scores (x-axis) & HAVCR2 vs. Mes (y-axis) (right). ‘N’ indicates the number of datasets that lie in the respective quadrant.

The correlation of these genes with epithelial and mesenchymal signatures revealed that *CD276* and *CD47* correlated positively with both Epi and Mes scores, although the association was stronger with the mesenchymal score as compared to the epithelial score **(Fig 5B, S5A, B)**. Expression of *HAVCR2* showed a positive correlation with the Mes score (16 vs 6 as in **Fig 5B**) but negatively with the Epi score (14 vs 7 as in **Fig S5C**). An antagonistic trend was seen in case of *LGALS9* which correlated primarily positively with epithelial signature (19 vs 4) but negatively with mesenchymal one (11 vs 9) (**FigS5D, 5B**). Thus, similar to PD-L1, additional immune checkpoint markers also seem to associate with a partial EMT signature.

Next, we compared the pairwise association of these four immune checkpoint markers with epithelial and mesenchymal scores. The predominant trends observed for all the genes was that the expression levels correlated positively with both epithelial and mesenchymal programs, with some checkpoints (except LGALS9) correlating positively with the mesenchymal program and correlating negatively with epithelial scores. For *LGALS9*, the opposite trend was observed i.e., *LGALS9* gene expression correlated negatively with epithelial scores but positively with mesenchymal ones (**Fig 5C**), supporting an association between partial or full EMT and upregulation of these immune checkpoints in oncovirus-positive samples.

## Discussion

Oncoviruses have been associated with multiple cancer hallmarks (Mesri et al. 2014), including metastasis and chemoresistance, and the formation of polyploid giant cancer cells (Chen et al. 2019; Herbein and Nehme 2020). Detailed molecular investigations into the oncovirus-driven alterations in cellular behavior can be instrumental in decoding how oncoviruses impact multiple stages of cancer progression and may pinpoint novel therapeutic strategies (Nečasová et al. 2022).

Here, we performed a meta-analysis of oncovirus-positive samples in interconnected axes of cellular plasticity: EMT, metabolic reprogramming, and immune checkpoint expression (Dongre et al. 2021; Jia et al. 2021; Loo et al. 2021; Lv et al. 2021; Sahoo et al. 2021). While crosstalk among these axes has been investigated in multiple cancers, their interconnection specifically in oncovirus-positive scenarios remains poorly understood. Our analyses identified consistent correlations between EMT, metabolism and PD-L1 levels across oncoviruses using bulk transcriptomics data. In particular, these analyses revealed that FAO and OXPHOS pathway activity is often decreased in samples with a strong enrichment of EMT signatures, implying a common adaptative mechanism connecting EMT and metabolic reprogramming. Despite identifying this consistent relationship, we were not able to pinpoint specific molecules mediating this interconnection or dissect how EMT and FAO (or OXPHOS) may influence one another. Intriguingly, glycolysis was higher in samples expressing both epithelial and mesenchymal genes, suggesting that glycolysis is likely to be associated with a hybrid E/M phenotype. This observation is reminiscent of the increase in glycolysis observed upon treatment with TGFβ, a canonical EMT inducer (Hua et al. 2020). Given that most studies focus on bulk-level analysis at limited time-points, it is challenging to convincingly demonstrate an association between hybrid E/M phenotype(s) and glycolysis; however, developing novel lineage tracing/barcoding strategies, coupled with single cell metabolomics, may pave the way for further investigation into the dynamics of cellular plasticity at different spatiotemporal coordinates in cancer evolution (Wei et al. 2022).

We also observed scenarios where immune checkpoints, such as PD-L1, correlated negatively with both OXPHOS and glycolysis, indicating that these axes of metabolic reprogramming are not strictly antagonistic to one another. These indications are strengthened by observations of high glycolysis /high OXPHOS and low glycolysis/low OXPHOS phenotypes, in addition to canonical high glycolysis/ low OXPHOS and low glycolysis/high OXPHOS ones (Yu et al. 2017; Jia et al. 2020; Amemiya and Yamaguchi 2022). However, in contrast to our analyses, glycolysis has been reported to enhance PD-L1 levels in cancer cells (Jiang et al. 2019). Further investigations shall be needed to better understand the context-specific mechanistic underpinnings and between immune-evasion and metabolic plasticity. Overall, our results show consistent trends in terms of partial EMT, metabolic plasticity, and immune checkpoint expression in oncovirus-positive samples.

## Author contributions

MKJ conceived research; MKJ and JAS supervised research; MS, RR, JMV and SK performed research. All authors contributed to writing and editing of the manuscript.

## Supplementary Figures

**Figure S1.**
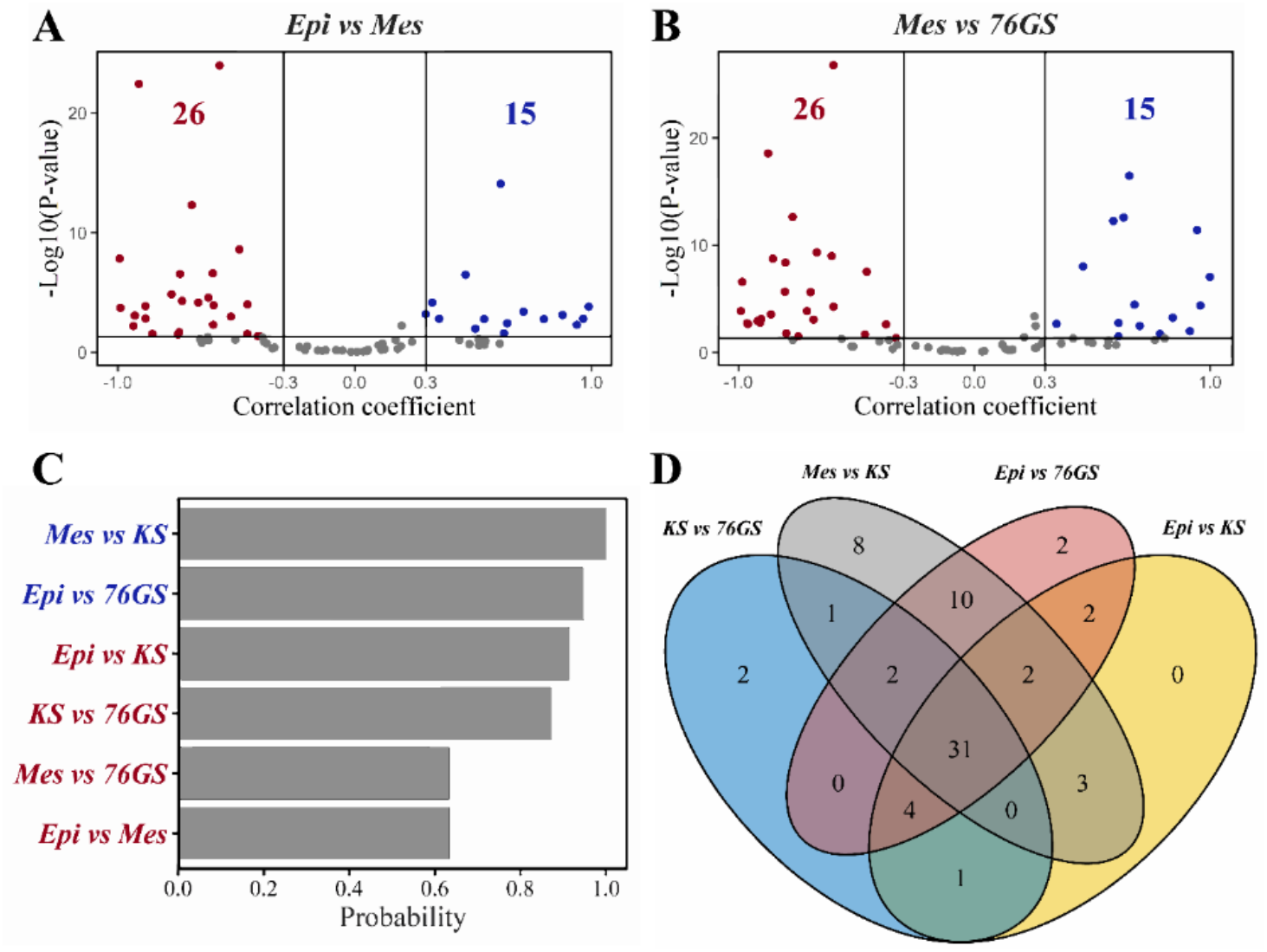
EMT Scoring metrics consistently report E/M status of samples across datasets. **A)** Volcano plot illustrating the Pearson correlation coefficient (x-axis) and the −log10(p-value) (y-axis) for Epi vs. Mes scores. Vertical boundaries are set at correlation coefficients corresponding to 0.3 and -0.3 and the cut-off for significant correlation is set at p<0.05. **B)** Same as A) but for Mes vs. 76GS scores. **C)** Probability of a dataset with significant (p<0.05) positive (blue) or negative (red) correlation for all possible pairs of E/M scoring metrics used in this analysis. **D)** 4-way Venn diagram for KS vs. 76GS scores, Mes vs. KS scores, Epi vs. 76GS scores, and Epi vs. KS scores representing datasets with significant correlation between the different EMT scoring metrics.

**Figure S2:**
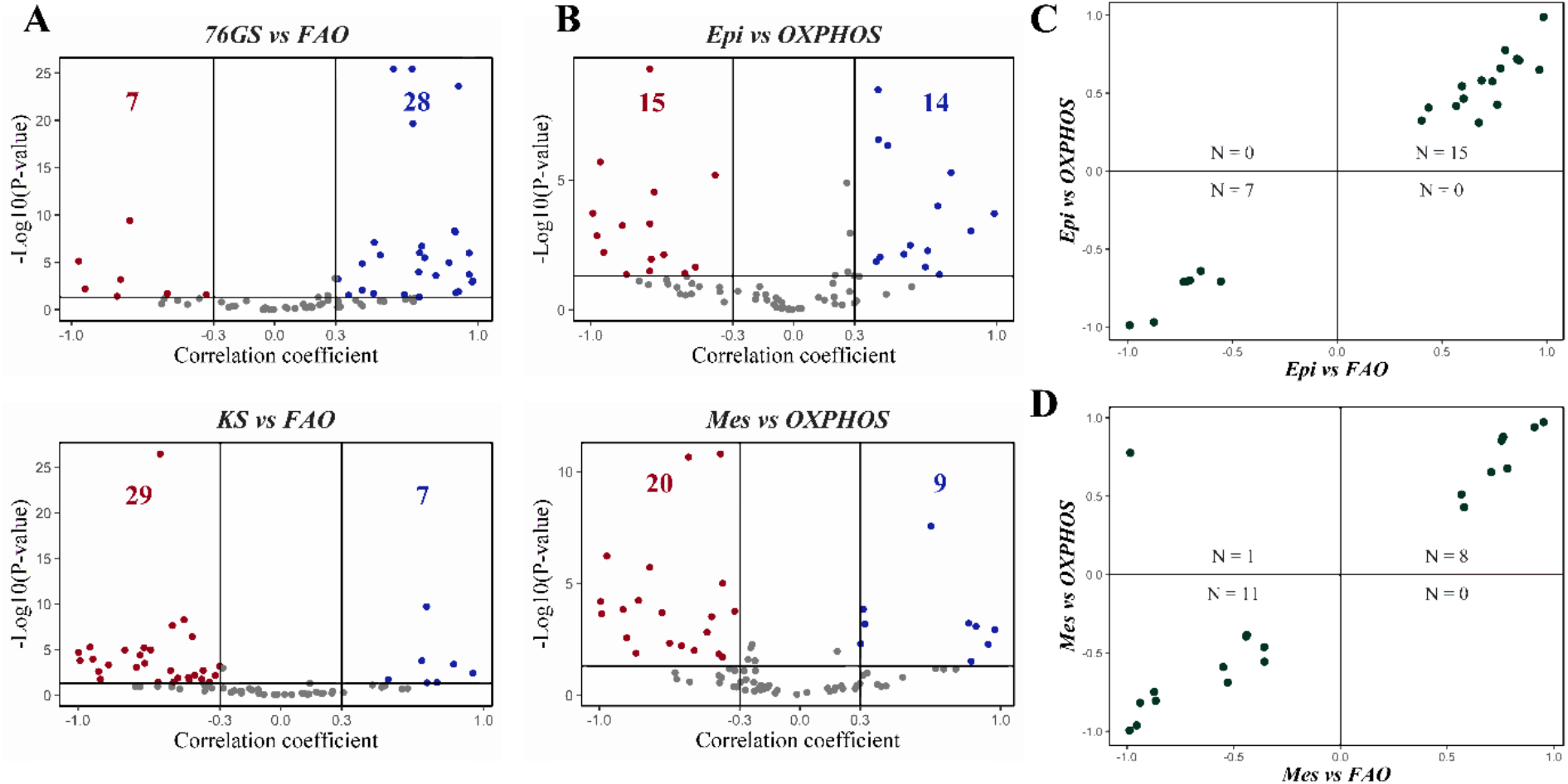
Association of EMT with OXPHOS and FAO programs. **A)** Volcano plot illustrating the Pearson correlation coefficient (x-axis) and the −log10(p-value) (y-axis) for 76GS vs. FAO (top) and KS vs. FAO scores (bottom). Vertical boundaries are set at correlation coefficients corresponding to 0.3 and -0.3, and the cut-off for significant correlation is set at p<0.05. **B)** Same as A) but for Epi vs. OXPHOS scores (top) and Mes vs. OXPHOS scores (bottom). **C)** 2D scatter plot illustrating correlation coefficients of Epi vs. FAO (x-axis) and Epi vs. OXPHOS scores (y-axis). ‘N’ indicates the number of datasets that lie in the respective quadrant. **D)** Same as C) but for Mes vs. FAO (x-axis) and Mes vs. OXPHOS scores (y-axis).

**Figure S3:**
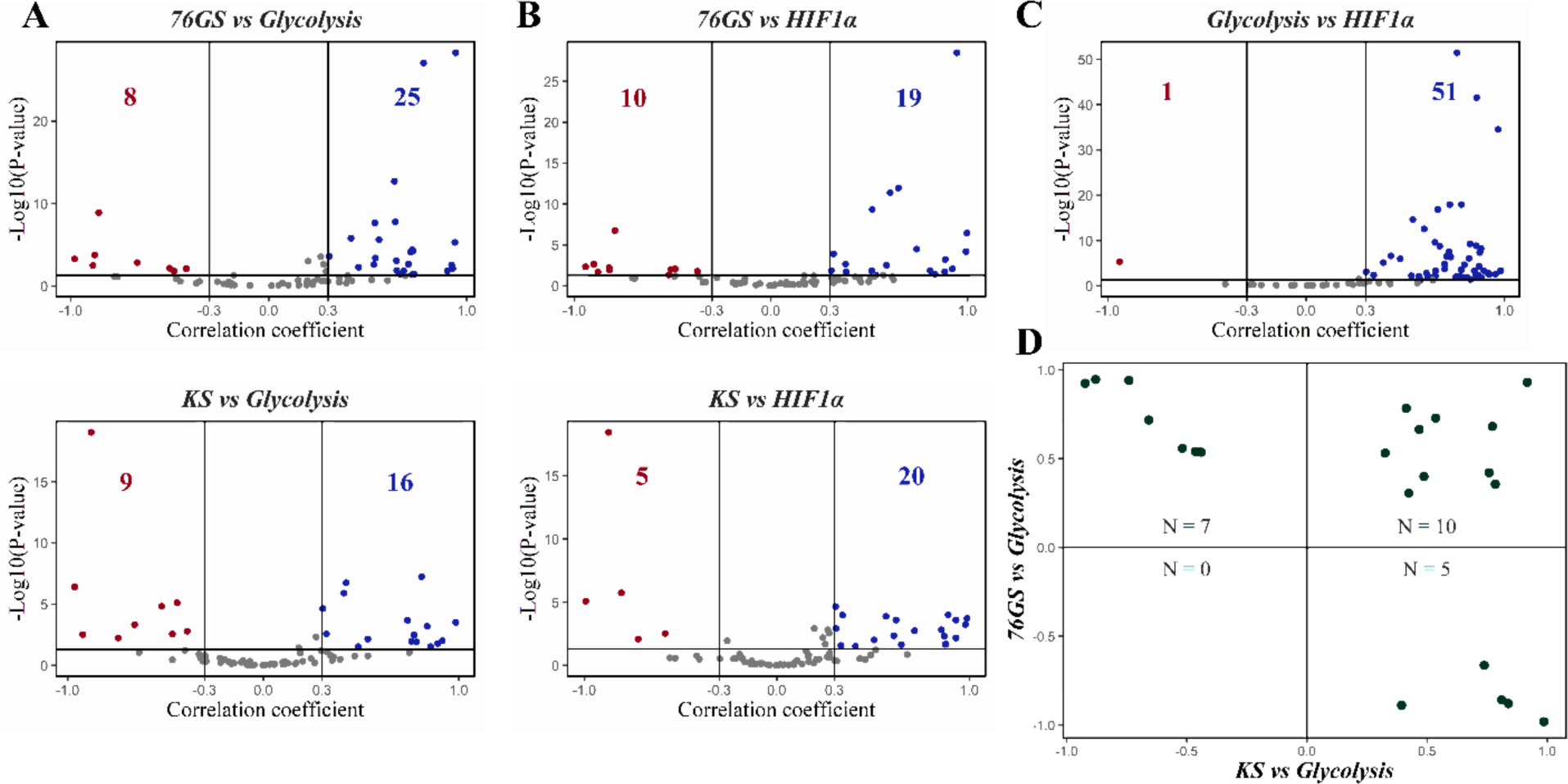
Association of glycolysis gene signature and its major driver-HIF1α. **A)** Volcano plot illustrating the Pearson correlation coefficient (x-axis) and the −log10(p-value) (y-axis) for 76GS vs. Glycolysis scores (top) and KS vs. Glycolysis scores (bottom). Vertical boundaries are set at correlation coefficients corresponding to 0.3 and -0.3, and the cut-off for significant correlation is set at p<0.05. Same as A) but for **B)** 76GS vs. HIF1α (top) and KS vs. HIF1α scores (bottom) and **C)** Glycolysis vs. HIF1α scores. **D)** 2D scatter plot of correlation coefficients of KS vs. Glycolysis scores (x-axis) and 76GS vs. Glycolysis (y-axis). ‘N’ indicates the number of datasets in respective quadrant.

**Figure S4:**
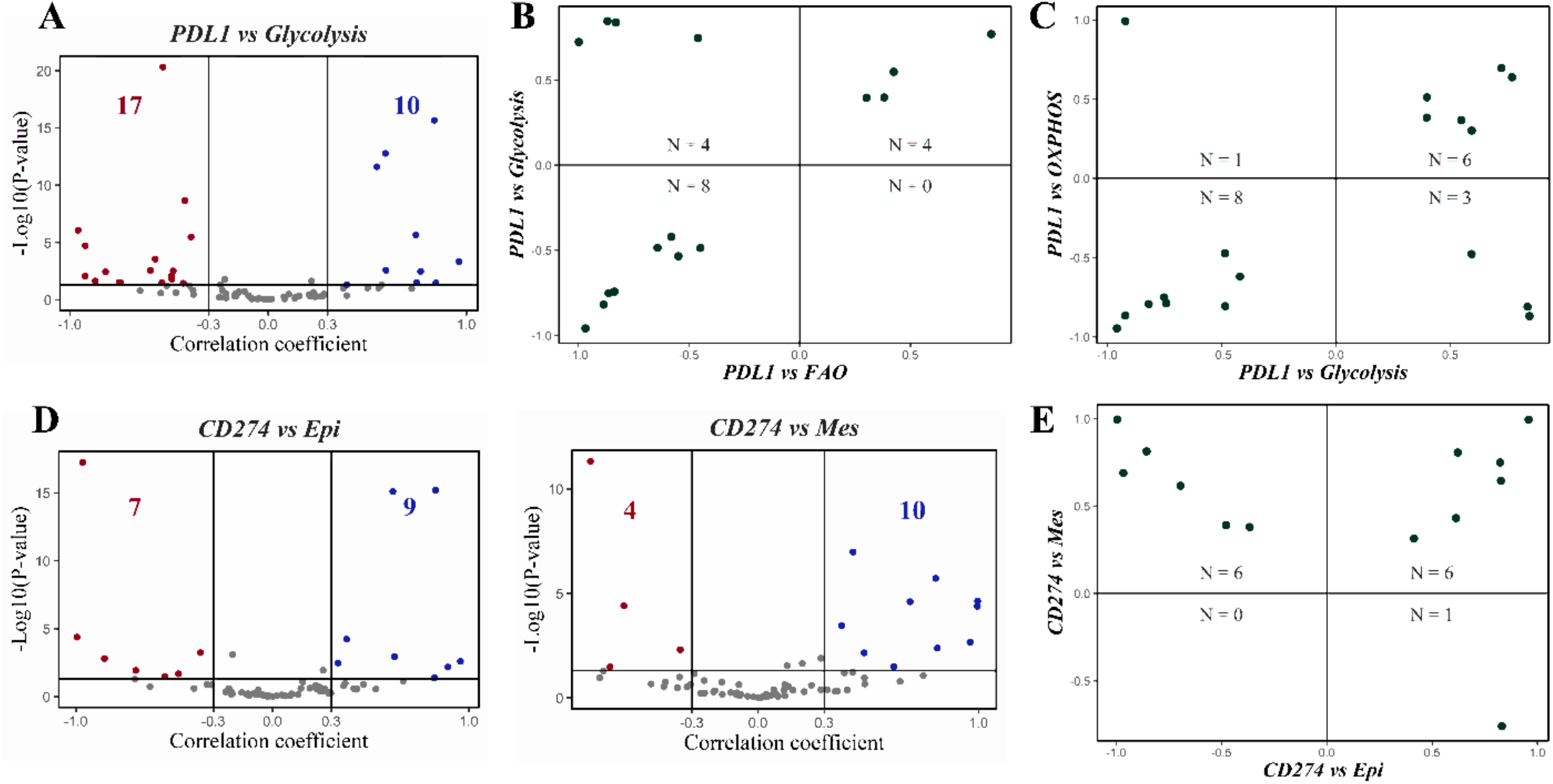
Different modalities of association between PD-L1 gene signature and CD274 gene expression with metabolic axes and EMT. **A)** Volcano plot illustrating the Pearson correlation coefficient (x-axis) and the −log10(p-value) (y-axis) for PD-L1 vs. Glycolysis scores. Vertical boundaries are set at correlation coefficients corresponding to 0.3 and -0.3, and the cut-off for significant correlation is set at p<0.05. **B)** 2D scatter plot of correlation coefficients of PD-L1 vs. FAO (x-axis) and PD-L1 vs. Glycolysis scores (y-axis). ‘N’ indicates the number of datasets that lie in the respective quadrant. **C)** Same as B) but for PD-L1 vs. OXPHOS (x-axis) and PD-L1 vs. Glycolysis scores (y-axis). **D)** Same as A) but for CD274 vs. Epi scores (left) and CD274 vs. Mes (right). **E)** Same as B) but for CD274 vs. Epi (x-axis) and CD274 vs. Mes scores (y-axis).

**Figure S5:**
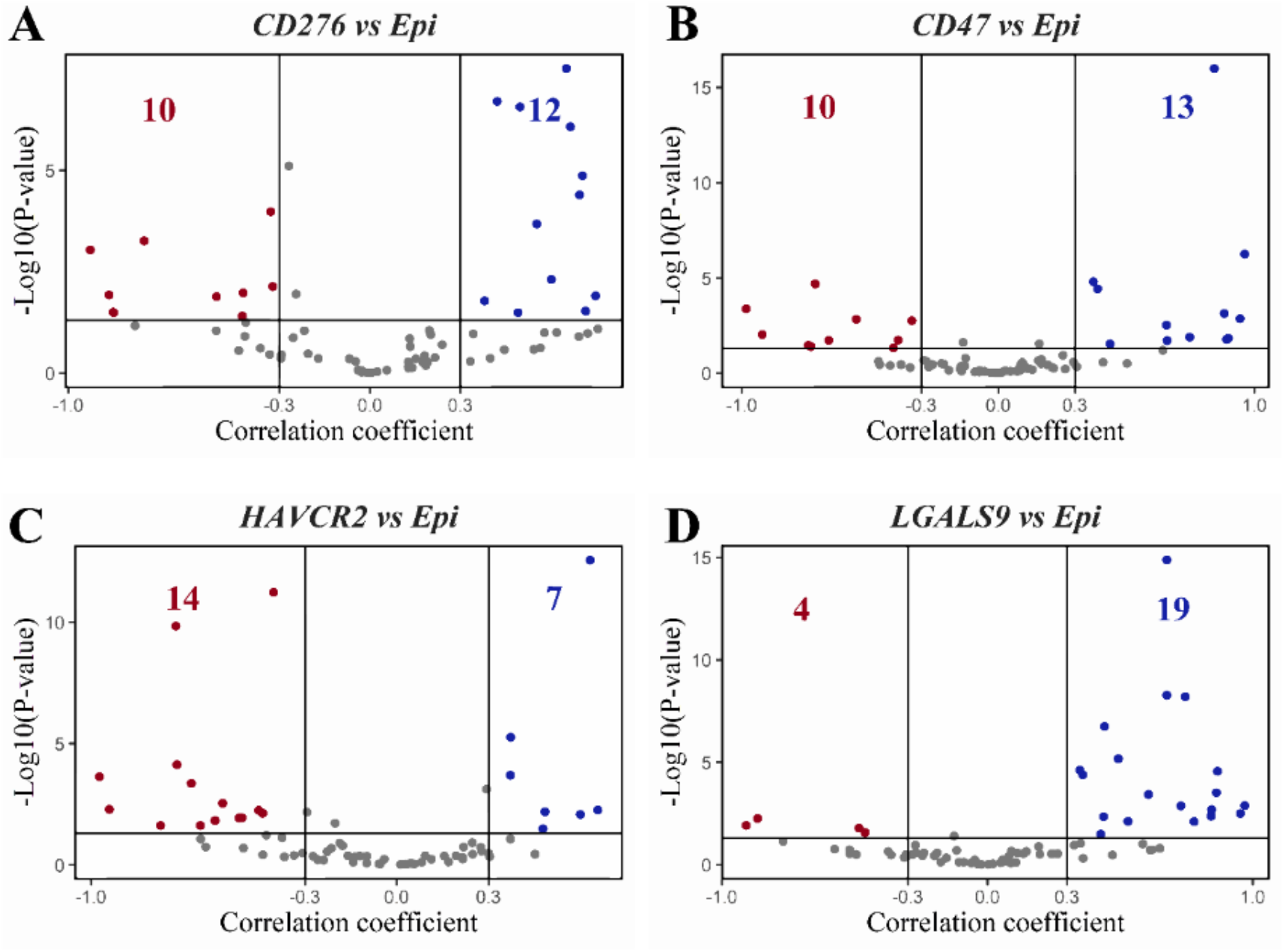
Association of different immune checkpoints with an epithelial program. **A)** Volcano plot illustrating the Pearson correlation coefficient (x-axis) and the −log10(p-value) (y-axis) for CD276 vs. Epi scores. Vertical boundaries are set at correlation coefficients corresponding to 0.3 and -0.3, and the cut-off for significant correlation is set at p<0.05. Same as A) but for **B)** CD47 vs. Epi, **C)** HAVCR2 vs. Epi and **D)** LGALS9 vs. Epi scores.

